# Stochastic Assembly Produces Heterogeneous Communities in the *C. elegans* Intestine

**DOI:** 10.1101/067173

**Authors:** Nicole M. Vega, Jeff Gore

## Abstract

**Author Summary:** Host-associated bacterial communities vary extensively between individuals, but it can be very difficult to determine the sources of this variation. In this manuscript, we demonstrate experimentally how randomness in colonization can result in large differences in the composition of host-associated bacterial communities, using the nematode worm *C. elegans* as a tractable host model. Here, the amount of variation between individual communities is a function of two rates relevant to how bacteria colonize the host intestine, the colonization rate and the birth rate. We can therefore control the degree of variation between communities by controlling the colonization rate, using the amount of bacteria presented to the worms to control the rate at which migrants enter the intestine. When worms are fed with two neutrally-competing fluorescently labeled bacterial strains at low colonization rates, we are able to produce noise-induced bistability in this system, where each community is dominated by bacteria of only one color. These results demonstrate the potential importance of noise as a driver of variation between communities and highlight the utility of the simple model organism *C. elegans* for studying questions relevant to host-associated microbial communities.

**Abstract:** Host-associated bacterial communities vary extensively between individuals, but it can be very difficult to determine the sources of this heterogeneity. Here we demonstrate that stochastic bacterial community assembly in the *C. elegans* intestine is sufficient to produce strong inter-worm heterogeneity in community composition. When worms are fed with two neutrally-competing fluorescently labeled bacterial strains, we observe stochastically-driven bimodality in community composition, where approximately half of the worms are dominated by each bacterial strain. A simple model incorporating stochastic colonization suggests that heterogeneity between worms is driven by the low rate at which bacteria successfully establish new intestinal colonies. We can increase this rate experimentally by feeding worms at high bacterial density; in these conditions the bimodality disappears. These results demonstrate that demographic noise is a potentially important driver of diversity in bacterial community formation and suggest a role for *C. elegans* as a model system for ecology of host-associated communities.

## Introduction

The gut microbiome varies greatly between individuals, and this variation could have important health consequences [1,2]. These differences may be due to deterministic differences such as genetic differences between individuals or differences in individual history and environmental exposure; stochasticity may also play a role in variation between individual communities [3,4]. Extensive research has focused on the deterministic factors directing microbial community composition [5–7]. Indeed, the canonical understanding of community assembly is based on deterministic niche-based processes, which push communities toward equilibria determined by the species present and the resources available. Under this view, if individual hosts are exposed to similar species and contain similar environmental conditions, they are expected to converge to similar community compositions. If there is heterogeneity between individuals, it comes from the presence of multiple equilibria in the system. Here, if an individual is perturbed away from its current stable condition, it will return to that condition; if the perturbation is large, it may instead converge to a new stable state.

By contrast, where stochastic processes dominate, similar hosts exposed to identical environmental conditions can diverge considerably in their final community composition. When heterogeneity between individuals is due to stochastic processes, individuals may exist in multiple states due to stochastic noise, but converge to a single deterministic equilibrium when the stochastic noise is small. This will be the case where migratory and other random processes outweigh any deterministic differences between colonizing species, such as competitive ability. While the stochastic processes of migration and drift are known to be important in determining the composition of natural communities [8,9], it is generally difficult to experimentally quantify the effects of stochasticity on community formation.

This is of particular interest in host-associated microbial communities, where a great deal of variation can be observed between individuals, and the role of stochastic processes in producing and maintaining this heterogeneity is not well understood. Despite substantial evidence for a core intestinal microbiota in multiple organisms [1,10–12], an increasing body of evidence suggests that stochastic processes may be generally important in determining the structure of host-associated communities [13–15]. Further, recent work suggests that a large amount of variation in the composition of these communities can potentially be explained by stochastic processes [16].

*C. elegans* is used in these experiments as a tractable experimental model of a gastrointestinal system undergoing bacterial colonization [17]. The worm is a bacteriovore, and during feeding, the intestine is colonized by the small fraction of bacteria which survive ingestion; these bacteria can survive and proliferate, forming communities within the host gut. The nematode has been used extensively as a laboratory organism and has many desirable properties, including selfing hermaphroditism (allowing maintenance of homozygous cultures), transparency under light microscopy, ease of culture, a short life cycle, and rapid generation time. Using the worm, it is therefore possible to generate large numbers of nearly genetically identical individuals with identical life histories, which can be colonized under identical conditions to allow observation of the stochastic forces underlying microbial community assembly. The simplicity of this system therefore allows a high level of control of both host and environment and a high degree of precision in measurement, both of which will be advantageous in observing the behavior of stochastically-assembled populations in the host environment.

## Results

To explore the role of stochastic colonization of an intestinal environment, we colonized *C. elegans* by allowing synchronized adult worms to feed for eight days on a 50/50 mixture of live *E. coli* fluorescently labeled with YFP or dsRed (**Fig. 1A**). Worms were colonized in liquid culture to ensure that all worms experienced a uniform environment without opportunities for selective feeding. Most worm intestines were successfully colonized by the bacteria, with an average of ~35,000 colony forming units (~3.82 ± 1.03 log_10_(CFU)) per worm (N = 53 colonized/56 total worms, mean ± s.d. of log-transformed non-zero counts). As expected, dilution plating of disrupted worm communities confirmed that the average community composition across all worms was close to 50/50 for the two colors (average 58% dsRed, **Fig. 1B**).

**Fig. 1.**
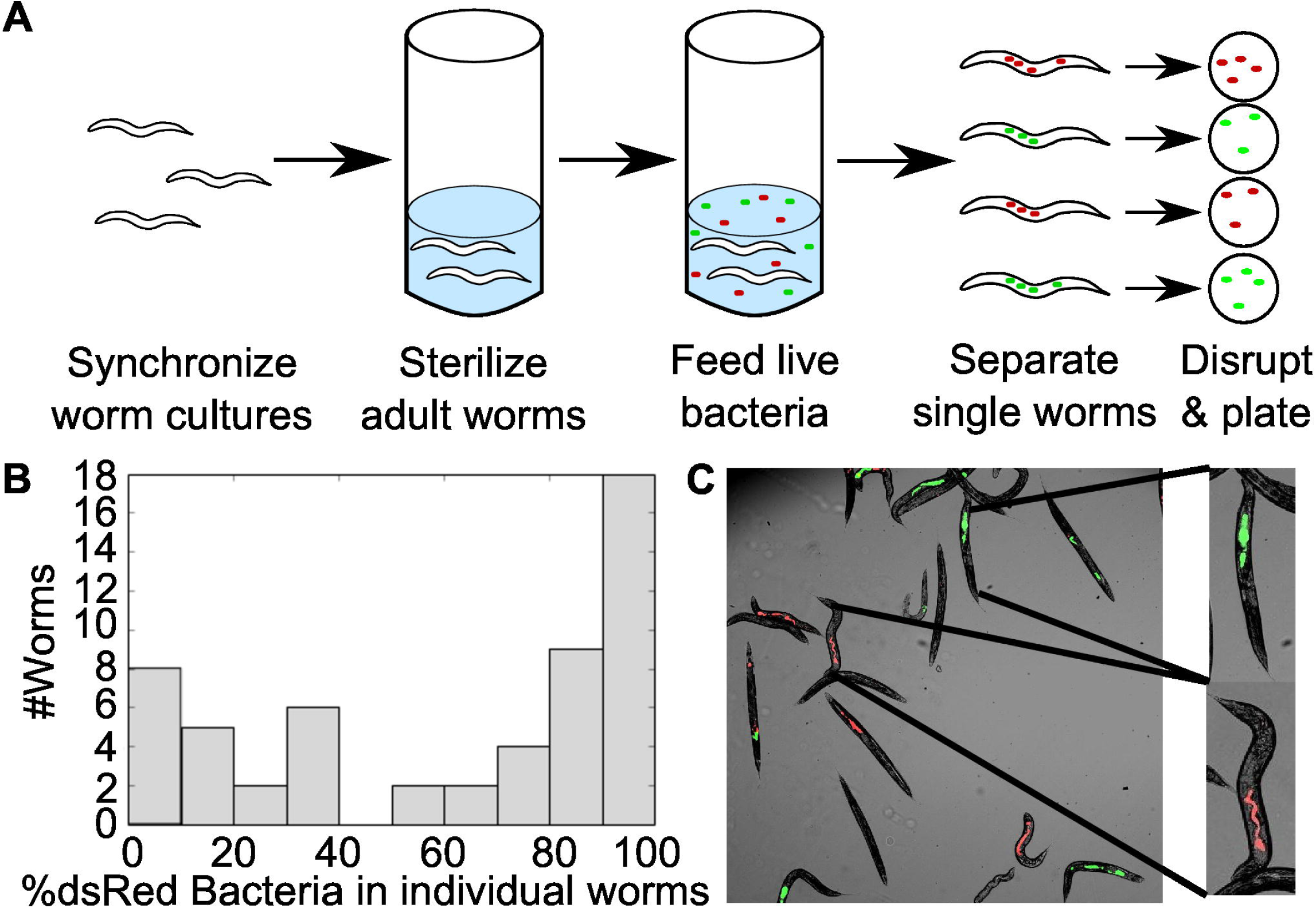
**Stochastic colonization produces bimodal community composition in the *C. elegans* intestine.** (A) Illustration of experimental procedure. Briefly, synchronized adult worms with germ-free intestines were fed for eight days on a 50/50 mixture of live *E. coli* (10^6^ CFU/mL in liquid culture) labeled with dsRed or YFP, then individual worms were isolated and disrupted to obtain intestinal contents. (B) After eight days of colonization, we observed a bimodal distribution in bacterial community composition (pooled data from three independent experiments, n=56; see S1 Data). (c) Bimodal community composition can be seen in fluorescence microscopy (three-channel overlaid image).

Given the large population sizes within the worm intestines and the fact that the worms were fed well-mixed bacteria at equal ratios, it would be natural to assume that the individual intestinal communities would also show a 50/50 mixture of the two colors of bacteria. However, though the average across all worms was close to the expected ratio, dilution plating of the intestinal communities of individual worms revealed that each worm tended to be dominated by one of the two colors, resulting in a bimodal distribution of community compositions (**Fig. 1B,C**). The intestinal communities within individual worms therefore display a striking difference from the 50/50 mixture present within the bacterial population present outside of the worm.

In the absence of deterministic factors such as competitive differences or life history differences between hosts, we conclude that the striking variation between individuals results from stochasticity in the colonization of the intestine. Indeed, fluorescent images of the colonized worms (**Fig. 1C**) revealed a patchy distribution of the two bacteria strains in the intestine, where bacteria have established monochromatic “clumps” rather than well-mixed colonies, consistent with rare colonization events leading to distinct colonies of each color. To further explore this hypothesis, we measured consumption rates and found that ~50 bacteria were consumed per worm per hour at the bacterial density used in feeding (**Fig. 2A)**, meaning that each worm consumed ~10^4^ live bacteria over the course of the experiment; bimodality was observed despite the large number of potential colonists, consistent with prior observations that a small fraction of ingested bacteria survive to colonize the *C. elegans* intestine (18). Indeed, we found that less than 1 in 1000 of the bacteria consumed by the worm both survive the feeding and successfully adhere to the intestine (**Fig. 2A**), thus providing a potential explanation for the source of stochasticity that leads to bimodality in community composition.

**Fig. 2.**
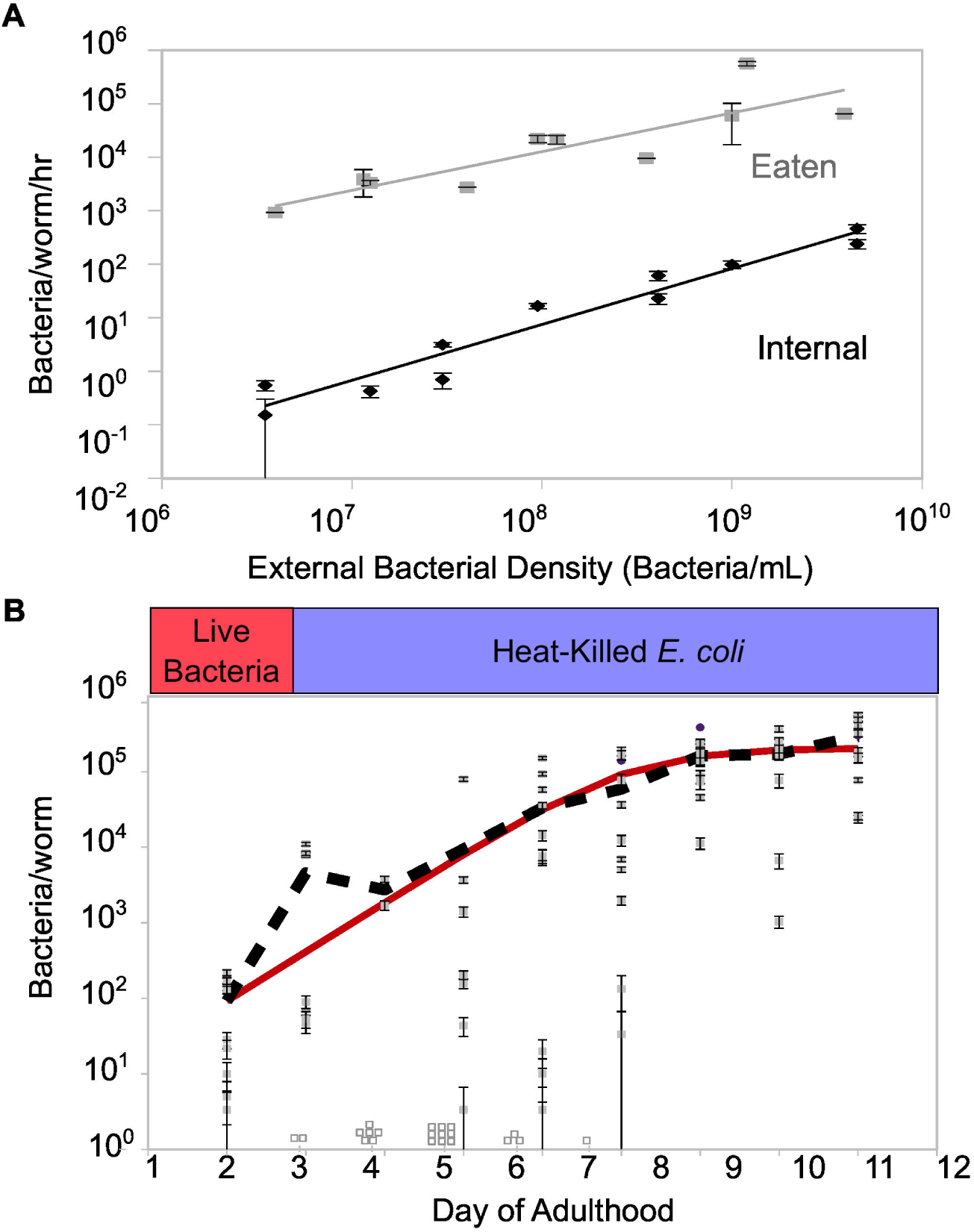
**Rare migrants grow and colonize the C. elegans intestine*C. elegans*intestine.** (A) External bacterial density during feeding controls feeding rate (Bacteria eaten/worm/hr, grey points and line), which in turn reflects the colonization rate (average number of internal bacteria acquired/worm/hr, black points and line). Worms were colonized in well-mixed liquid culture with *E. coli* MC4100 E3350 or E3322 as a food source; no differences were observed in colonization ability between strains at one hour, and data were pooled. Average internal CFU/worm measurements were obtained from batch digests of 100-200 worms after one hour of colonization. (B) In the absence of new colonization (see Methods), growth of unlabeled *E. coli* MC4100 in the *C. elegans* intestine is described by a logistic model (black line, average CFU/worm; red line, fit of the logistic equation with *r* ≈ 1.5 day^-1^ and *K* ≈ 2 x 10^5^ CFU/worm). Open squares indicate number of individual uncolonized worms digested each day (CFU/worm = 0). In both panels, bars indicate mean ± SD estimated by assuming Poisson error of colony counts. See S1 Data.

To gain insight into the process of stochastic colonization and growth, we constructed a minimal model which incorporates stochastic colonization of the worm at rate *c* (0.1-100 bacteria/hr), birth rate *b* (0.6 hr^-1^), saturated population size *K* (2 * 10^5^ bacteria/worm), and death rate *d* (0.54 hr^-1^; for discussion see Methods) which includes both death and excretion out of the worm (**Fig. 3A, Fig. S1**). The deterministic dynamics are represented using a logistic equation:

**Fig. 3.**
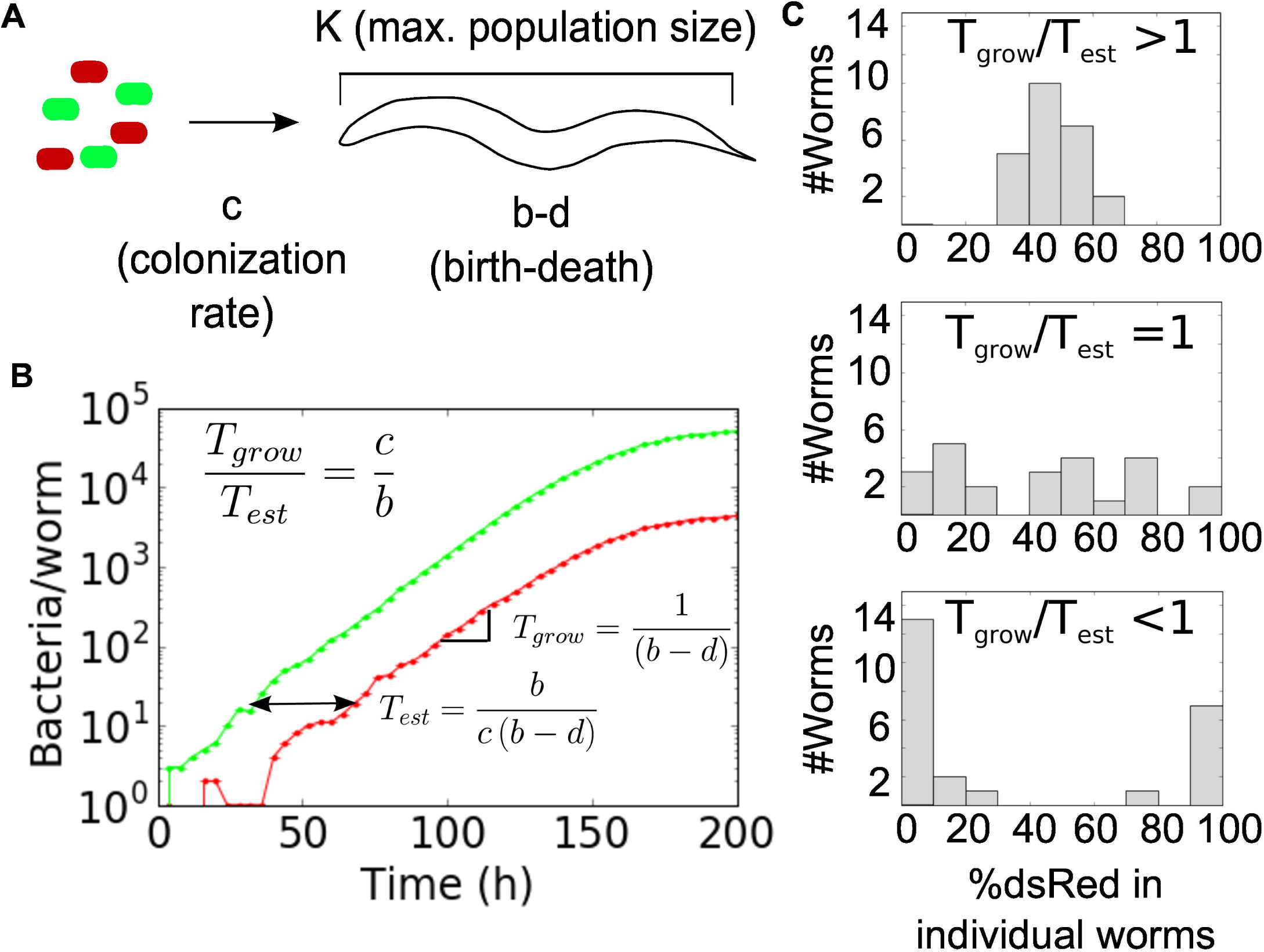
**A stochastic model describes colonization of the C. elegans intestine*C. elegans*intestine.** (A) Bacterial population dynamics in the worm were modeled using a density-dependent logistic framework, where worms are colonized at a rate *c*, bacteria grow and die within the worm at rates *b* and *d*, and populations within the worm saturate at a carrying capacity *K*. All parameters are constant in time and identical for both bacterial strains. (B) A single Gillespie stochastic simulation (GSSA) run using a low colonization rate (*c*=0.1 hr^-1^) is presented to illustrate the different timescales that determine community assembly in this system. In simulations, worms were colonized with a 50/50 mix of identical bacterial strains, shown here as green and red lines. *T*_*grow*_ is the characteristic timescale of colony growth inside the worm, and *T*_*est*_ is the expected time between successful colonization events. (C) Simulations were performed at a range of colonization rates (top panel, *c*=10; middle, *c*=*b*=0.6; bottom panel, *c*=0.1) to illustrate how the critical ratio *T*_*grow*_*/T*_*est*_ = c*/b* controls the transition from bimodal to unimodal community composition. See Methods, S2 Data, and S4 Data for code.

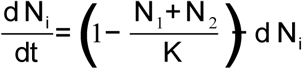

Since noise is important in driving the behavior of this system, we conduct simulations using the corresponding generalized system of reactions:

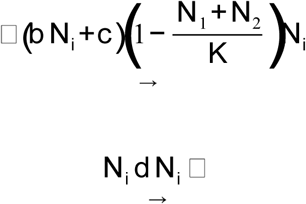

where *i*=1,2 denotes the index of the neutrally-competing bacterial strain. Strains are identical apart from their index and use the same values for *b, d, K,* and *c*. Stochastic simulations were implemented using the tau-leaping method [19,20]. Notably, for a simplified version of the birth-death-immigration process where we ignore saturation, the full time-dependent solution can be found analytically [21].

Colonization rate in this context corresponds to the rate at which bacteria survive being consumed and successfully adhere to the gut. We determined that growth of bacteria in the *C. elegans* intestine in the absence of colonization is highly variable, but average population size over time can be described by a logistic model (**Fig. 2B**), consistent with previous reports in individual zebrafish [22]. In this system, colonization rate (number of live intestinal bacteria obtained from the environment per unit time) is an experimentally tunable parameter which depends on the density of bacteria outside the worms (**Fig. 2A**).

In our stochastic model, we expect that the variation between communities in individual worms will depend on the relative magnitude of two time-scales: (1) the typical time between successive successful colony establishment events 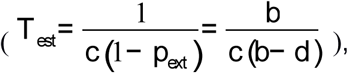 where 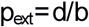 is the probability of a new colonist going extinct [23]) and (2) the timescale associated with net exponential growth of a colony once established 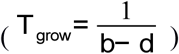 Fig. 3B,C). As can be seen from these equations, the establishment time *T_est_* is determined by the rate of colonization and the birth and death rates, since these together influence the probability that an initial migrant will successfully start a new colony within the worm. If 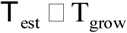, new colonists are expected to arrive rapidly relative to the time it takes a colony to grow; the distribution of bacteria within each intestine will be forced by the external bacterial population, resulting in worm-associated colonies that are very similar to both the pool of colonists and to one another. However, if 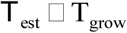, successful colonists are expected to be rare and a given colonist is expected to have a long time to grow and fill the host gut before another successful colonist arrives, resulting in worms that are dominated by the single-color offspring of their first successful colonist.

Which regime we are in therefore depends upon the ratio 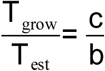. Biologically, birth rate *b* is fixed within a fairly narrow range of values (less than a full order of magnitude), capped by the maximum physiological growth rate of the bacteria and bounded below by the observed net growth rate of the population (given by b-d). However, the colonization rate *c* can be easily tuned over several orders of magnitude by changing the density of bacteria on which the worms are allowed to feed **(Fig. 2)**. The model thus predicts that if we increase the colonization rate *c* then the bimodality will disappear, and each intestinal community will approach a 50/50 mixture of the two labeled bacteria.

To test this hypothesis, we fed worms at a range of higher bacterial cell densities, thus increasing the colonization rate (**Fig. 4A**). As expected from the model, as bacterial concentrations outside the worm increase, the resulting intestinal communities become more similar to one another (**Fig. 4A**), and the distribution of populations goes from bimodal to unimodal (**Fig. 4B**). Specifically, at high bacterial density, each worm has a microbial community that is reflective of the 50/50 mixture of red/green bacteria present outside of the worm. (Corresponding bacterial abundance data are shown in **Fig. S2A**.) The strength of stochastic forces in determining the composition of the worm intestine therefore depends strongly on the feeding rate of the worm. Despite the fact that these bacteria compete neutrally in the absence of worms (**Fig. S3A-B**) and colonize worms at comparable rates (**Fig. S3C**), populations inside the worms show a slight excess of dsRed-labeled bacteria, most likely due to differences in growth and/or death rates inside the worm gut (**Fig. 4A, S3E-F**). Notably, we expect the observation of a bimodal to unimodal transition to be robust to the choice of experimental duration before harvesting (**Fig. S4**).

**Fig. 4.**
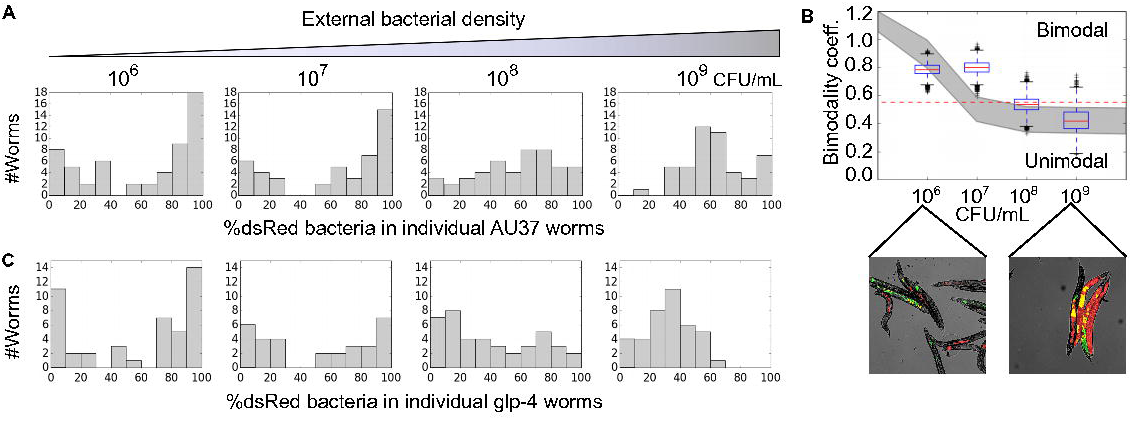
**Colonization rate tunes heterogeneity of microbial community composition between hosts.** (A) Adult AU37 worms fed for eight days on a 50/50 mixture of dsRed and YFP-labeled *E. coli* over a range of concentrations from 10^6^-10^9^ CFU/mL show a transition from bimodal to unimodal community composition. Data are pooled from 2-3 independent experiments (n=48-56 worms). (B) The transition to bimodality in (A) is confirmed by calculation of the bimodality coefficient (see Methods) for these data (bootstrap confidence intervals over 10,000 runs, red dashed line indicates BC_crit_=0.55) and by direct observation of intestinal communities via fluorescence microscopy. Grey area represents mean ± 1 SD of GSSA simulations exploring the effects of parameter uncertainty (K = (30000, 300000); b = (0.1, 0.6); d = (0.1, 0.5); b>d; see Methods) (C) The transition to bimodality is also observed in an immune-competent strain of *C. elegans* (*glp-4*, n=31-45 worms). This strain possesses the same temperature-sensitive sterile mutation as AU37 but lacks the mutation to immune system function (*sek-1*). See S1 Data.

A prediction of our model (**Fig. 3**) is that the transition from bimodality to unimodality does not depend upon the carrying capacity of the worm, since the bimodality arises from initial differences in colonization time that are “frozen in” to the final distribution without regard to final population size. To test this prediction, we take advantage of the fact that the population size of the gut community depends strongly on the functioning of the immune system of the worm. The experiments described so far were performed in an immune-compromised worm strain capable of supporting large bacterial populations (AU37), whereas a worm strain with increased immune function (glp-4) has a bacterial population size that is an order of magnitude lower (~4,700 bacteria per worm as compared to the 35,000 observed previously); in addition to the lower effective carrying capacity observed in glp-4 worms, death and colonization rates may also differ from those observed in AU37 due to the highly active immune system of the glp-4 strain. Consistent with predictions from the model, the transition from bimodality to unimodality also occurs in non-immunocompromised worms, and this transition occurs at a similar density of bacteria (**Fig. 4C, S5;** wild-type worm data shown in **Fig. S6**). Our experimental observation of heterogeneity in gut populations driven by stochastic colonization is therefore was robust to host genotype, even when the different hosts result in steady-state bacterial population sizes that differ by more than an order of magnitude.

To determine whether stochastic colonization also plays a role in shaping the outcomes of inter-species competition within the gut, we fed *C. elegans* on a 50/50 mixture of *Enterobacter aerogenes* and *Serratia marcescens*, both Gram-negative bacteria from the family *Enterobacteriaceae* (**Fig. 5A**). Unlike the single species case, the resulting communities were not distributed around the 50/50 mark, reflecting the more complex dynamics in this system where the two species do not compete neutrally. Under conditions where colonization is rapid (worms fed at ~10^10^ CFU/mL), the distribution of populations in individual worms had a visible central tendency at ~70% *E. aerogenes*, possibly reflecting the more rapid colonization displayed by this species at high feeding densities (**Fig. S7A**), and variation between individual worms was low (**Fig. 5B,C**). Under slow colonization conditions (worms fed at ~10^6^ CFU/mL), however, the median falls to 10-20% *E. aerogenes*, possibly due to a competitive advantage for *S. marcescens* inside the worm (**Fig. S7D,E)**, and the worm intestinal communities tend to be entirely dominated by one of the two species (**Fig. 5B,C**). Indeed, we find that a minority of worms are dominated by *E. aerogenes* despite the fact that the distribution is peaked at 10 – 20% *E. aerogenes*, highlighting that while interspecies competition will likely result in richer dynamics than are present with neutral labels, stochastic colonization is nonetheless important in shaping the heterogeneity between communities.

**Figure 5.**
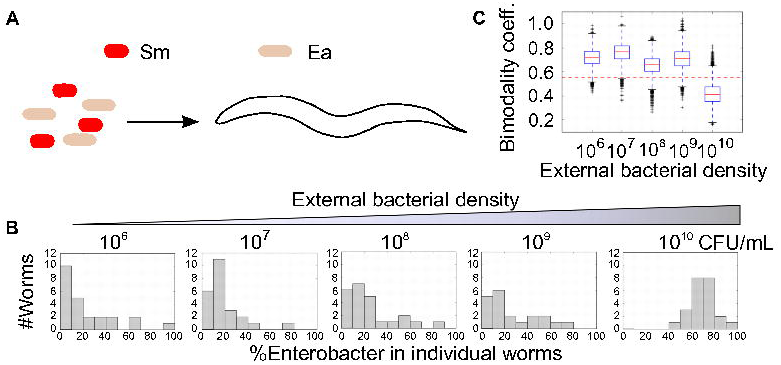
**Stochastic colonization shapes assembly of a simple multi-species community.** (A-B) The transition to bimodality was observed when adult AU37 worms were fed on a 50/50 mixture of *Serratia marcescens* and *Enterobacter aerogenes* at a range of colonization densities (10^6^-10^10^ CFU/mL) for five days (see S1 Data). The distribution of communities at the highest colonization density is significantly different than that observed at any lower density (Mann-Whitney U test, p=1.34*10^-6^ -3.33*10^-8^). (C) The transition to bimodality in %Ea is demonstrated (BC calculations for bootstrapped data using 10,000 repetitions, see S3 Data for code).

## Discussion

In this study we have found that stochastic colonization can lead to dramatic differences in the gut communities present between different hosts. Here we observe bimodality in community composition with neutral labels which is due to stochasticity in colonization. We found that this heterogeneity between worm hosts can be explained by a simple stochastic model, and that this model accurately predicts that our system will shift from a bimodal to a unimodal distribution of community compositions as colonization rate increases. Our results indicate that stochastic colonization is dominant at bacterial densities under 10^8^ CFU/mL; these densities are within the range expected in natural soil [24], and it is perhaps surprising that stochastic colonization can dominate at such large bacterial densities. However, the large degree of bottlenecking between the environment and the host intestine, here due in part to mortality caused by the worm grinder, provides the randomizing element necessary to explain these results. Bottlenecking of this magnitude may be relevant for intestinal microbial populations in other systems; interestingly, the degree of mortality observed in bacteria in these experiments is similar to that observed in transit of *V. cholerae* to the mouse gut [25]. It is important to note that the variation in community composition is not due to the number of colonists being trivially small; at bacterial densities of 10^8^ CFU/mL, each worm is exposed to ~10^3^ potential colonists over the course of the experiment (**Fig. 2A).** Rather, according to the predictions of our model, variation in community composition arises due to exponential growth of early successful colonists, which prevents the formation of large populations by later colonists.

Immune function of the host may also be expected to play a role in controlling colonization of the intestine. As we have seen, a host strain with high immune function supports a much smaller bacterial population than a corresponding immune-compromised strain. We do not expect the difference in carrying capacity to have a significant effect on the outcome of stochastic colonization. However, immune activity could potentially also alter the effective colonization rate by causing mortality of potential colonists after passage through the grinder. In this case, we would expect the transition from bimodality to unimodality to occur at a higher bacterial feeding density for a host with high immune function than for an immune-compromised host; a close examination of the data reveals some support for this hypothesis (**Fig. S5**), but the difference, if any, in transition point between worm strains in this study is small (**Fig. 4**).

Despite the explanatory power of the simple stochastic model, the experimental data do show more variation than predicted by the model. In particular, there is more variation in the number of bacteria per worm than can be explained using our simple model (**Fig, 4B, Fig. S2**). Further, the transition from bimodality to unimodality occurs at a somewhat higher bacterial feeding density than expected; specifically, at ~10^8^ CFU/mL rather than the 10^7^ CFU/mL that would be predicted by the model using our estimates of parameters such as the net colonization rate *c* and the birth rate *b* of bacteria within the gut (**Fig. 3C, 4**). The actual birth rate *b* could be lower than assumed in Fig 4C, but this would actually shift the transition to even lower bacterial densities as the transition occurs when *c = b*. Despite its utility in providing insight into the mechanism of noise-induced bimodality, this simple stochastic model therefore does not capture all of the variation present in our population of hosts.

This suggests that the simple model used here does not account for one or more important sources of variance. For example, despite the use of age-synchronized adult worms, there may be variation between host environments; that is, the demographic parameters directing community formation may vary among individual hosts. Indeed, considerable heterogeneity in “effective age” of individuals has been observed within synchronized cultures of *C. elegans*, resulting in heterogeneity in stress resistance and lifespan [26,27]. Similarly, these parameters may change in value over time (and therefore over the course of an experiment) for individual worms. In addition, the demographics of bacterial growth and death may change as worms age [18,28,29], and excretion from the worm may occur in discrete pulses consisting of many individuals, as recently observed in a zebrafish model [30]. These sources of variation could have profound effects on community composition, particularly in more complex cases where there are higher numbers of colonizing species and where the colonists do not compete neutrally, or where interactions between bacteria and host occur over evolutionary time [31].

The bimodal-to-unimodal transition with increasing colonization rate observed here resembles the expected behavior of a population behaving according to Hubbell's neutral model of island biogeography [32]. Unlike our model which begins with an uncolonized “island”, the Hubbell neutral model begins with an island of size *N* which is filled to capacity, in which individuals cannot be displaced but dead individuals can be replaced through either migration from a metacommunity or reproduction from within the island. In this zero-sum model, what determines whether the population distribution is unimodal or bimodal depends not on T_grow_ but rather on the typical timescale over which drift leads to extinction of a neutral lineage (which is much longer than T_grow_ and scales as the square of the population size *N*). The bimodality that we observe experimentally may therefore in principle be transient, but it is effectively the equilibrium distribution since the transient would last longer than the lifetime of the worm.

The work presented here suggests potential for *C. elegans* as a model organism for host-associated microbiotal community formation. The worm is used here as a tractable experimental model of a gastrointestinal system undergoing bacterial colonization [17]. The worm has a complex relationship with bacteria: it is a bacterial predator [33] which can be injured by pathogens and by overgrowth of “innocuous” bacteria within its intestine [18], while benefiting from probiotic effects and small molecule production by apparent “symbionts” [34,35]. The worm intestine can be colonized with a wide variety of bacterial species, including important human pathogens [36–38]; these bacteria grow and interact within the host, and their interactions are shaped by the properties of the host environment [36]. *C. elegans* is a selective environment, allowing some bacterial species to thrive while preventing the growth of others, and there is some evidence for a potentially beneficial “core” gastrointestinal microbiota in the worm [38–40]; in future studies, it would be interesting to explore the heterogeneity between individual worm-associated communities formed under more natural conditions from a rich, realistic metacommunity and to evaluate the effects of stochastic colonization under these conditions [9,16]. Though we did not observe any difference in heterogeneity between individual worms when colonizing immune-impaired worms vs. worms with active immunity with a single bacterial species, there is considerable evidence that the *C. elegans* innate immune response differs depending on the microbial species to which the worm is exposed [41–44]; it is plausible that the immune response could play an active role in shaping the composition of a true multispecies community in this host. Though the direct interactions between *C. elegans* and its naturally associated intestinal communities are not well understood, there is evidence that the worm can benefit directly from the presence of live bacteria [17,34,45,46]. In brief, *C. elegans* provides a well-controlled, biologically interesting, highly replicable environment for microbiotal experiments, and the work shown here demonstrates potential utility for future studies of stochastic and deterministic processes in community assembly.

The results presented here suggest that stochastic processes may have strong effects on host-associated microbial community assembly. Stochasticity has long been understood as a potential source of variation in these communities, but much work in this area focuses on deterministic factors such as differences between individual hosts. However, stochasticity is known to be a very important source of variation in some cases; for example, clonal heterogeneity between hosts has been observed in infectious disease, where an individual host may be dominated by a small number of strains or even a single clonal lineage, and stochastic bottlenecking during the life cycle of the disease has been implicated as a cause of this heterogeneity [47–49]. Due to the effects of bottlenecking, stochastic assembly may therefore be an important factor in evolution within host-associated bacterial communities and in co-evolution of bacteria and host [50]. Future studies will be necessary to determine the degree to which stochastic colonization drives heterogeneity in the microbiota in other animals.

## Methods

### Nematode culture

Unless otherwise stated, nematodes were cultivated under standard conditions [51], and the temperature-sensitive reproductively sterile *C. elegans* strain AU37 (*glp-4(bn2)* I; *sek-1(km4)* X) [52] was used for experiments. Due to the *glp-4* mutation, this strain is able to reproduce at 15°C but does not develop gonads and is therefore sterile when raised at room temperature (23-25°C); use of this strain prevented the worms from producing progeny during experiments, ensuring that all worms in a given experiment were of the same age and had the same life history. Additionally, the loss of the large hermaphrodite ovary was advantageous for microscopy due to the simplified internal tube-within-a-tube body structure. The *sek-1* mutation decreases immune function, making these worms more susceptible to bacterial colonization and allowing large intestinal bacterial communities to be readily established [52]. Worm strains were provided by the Caenorhabditis Genetic Center, which is funded by NIH Office of Research Infrastructure Programs (P40 OD010440).

Synchronized cultures were obtained using standard protocols [51]. For propagation of worm cultures, AU37 cultures were maintained at the permissive temperature (15°C) on NGM agar plates with lawns of the standard food organism *E. coli* OP50. N2 worms were maintained at 23°C on NGM plates with OP50. For synchronization, we washed 4-6 nearly starved plates with sterile distilled water and treated the harvested worms with a bleach-sodium hydroxide solution; the isolated eggs were placed in M9 worm buffer overnight to hatch, then transferred to NGM + OP50 plates at the non-permissive temperature (25°C) for 3 days to obtain gonadless (sterile) synchronized adults. Adults were washed from plates using M9 worm buffer + 0.1% Triton X-100, then rinsed with M9 worm buffer and surface bleached for 10 minutes at 4°C in a 1:1000 bleach solution in M9 to remove any live OP50. Worms were then transferred to S medium + 100 μg/mL gentamycin + 5X heat-killed OP50 for 24 hours to kill any OP50 inhabiting the intestine, resulting in germ-free synchronized worms. These worms (2-3 day synchronized adults) were then rinsed in M9 worm buffer + 0.1% Triton X-100, washed via sucrose flotation to remove debris, and rinsed 3X in M9 worm buffer to remove sucrose before use in experiments.

### Bacteria

These experiments used *E. coli* MC4100 (F^-^ [araD139]B/r Δ(argF−lac)169* &lambda^−^ e14-flhD5301 Δ(fruK-yeiR)725 (fruA25)‡ relA1 rpsL150(strR) rbsR22 Δ(fimB-fimE)632(: IS*1*) *deoC1*) carrying plasmids E3350 (pUC18T-mini-Tn7-Gm-eyfp, accession DQ493879) or E3322 (pUC18T-mini-Tn7-Gm-DsRedExpress, accession DQ493880)[53]. *E. coli* MC4100 was obtained from the *E. coli* Genetic Stock Center (CGSC #6152). For multispecies colonization experiments, *Enterobacter aerogenes* (ATCC 13048) and *Serratia marcescens* (ATCC 13880) were obtained from ATCC.

Bacterial strains were grown in individual cultures in LB + 30 μg/mL gentamycin for selection where necessary. For feeding assays, *E. coli* were grown overnight at 37°C, and *E. aerogenes* and *S. marcescens* were grown overnight at 30°C. Bacterial cultures were acclimated briefly (~1 hour) to room temperature before feeding. To construct feeding cultures, *E. coli* were centrifuged 1 minute at 9K RPM to pellet, washed once in S medium, then resuspended in S medium + 30 μg/mL gentamycin. As *S. marcescens* is highly motile and very difficult to pellet through centrifugation, *E. aerogenes* and *S. marcescens* were chilled for 30-60 minutes before centrifugation, then centrifuged 1 minute and 10K RPM to pellet before washing and resuspension.

### Colonization of *C. elegans*

Sterile, washed adult worms were resuspended in S medium (with 30 μg/mL gentamycin for selection where needed) and moved in 900 μL aliquots to individual wells of a 24-well culture plate, with a final concentration of ~1000 worms/mL. 100 μL of bacterial suspension at 10X desired concentration was added to each well before plates were covered with an adhesive gas-permeable microplate sealing film (USA Scientific). Plates were incubated with shaking at 300 RPM at 25°C. For multi-day assays, worms were re-fed every 24 hours; to do this, worms were removed from wells, washed 1X with M9 worm buffer + 0.1% Triton X-100 and 2X with M9 worm buffer to remove most external bacteria, and resuspended in 900 μL fresh S medium for re-plating as previously described.

### Disruption of worms and plating of intestinal populations

Colonized worms were washed 1X with M9 worm buffer + 0.1% Triton X-100 and 2X with M9 worm buffer to remove most external bacteria, then resuspended in 100 μL S medium + 1X heat-killed OP50 and incubated at room temperature (23°C) for 30-60 minutes to allow the worm to purge any non-adhered bacterial cells from the intestine. Worms were then rinsed 2X with M9 worm buffer + 0.1% Triton X-100, chilled briefly to stop peristalsis, and surface bleached for 10 minutes at 4C in 100 μL M9 worm buffer with commercial bleach at 1:1000 concentration.

For manual disruption, bleached worms were rinsed 1X in M9 worm buffer + 0.1% Triton X-100 to remove bleach, then transferred to 3 mL M9 worm buffer + 1% Triton X-100 in a small (40 cm) petri dish (Fisher Scientific). Individual worms were pipetted out in 20 μL aliquots of buffer and transferred to 0.5 mL clear microtubes (Kimble Kontes) for manual disruption with a motorized pestle (Kimble Kontes Pellet Pestle with blue disposable pestle tips, Fisher Scientific). Magnification was provided by a magnifying visor (Magni-Focuser Hands Free Binocular Magnifier 3.5X). After disruption, tubes were centrifuged 2 minutes at 12K RPM to collect all material, and the resulting pellet was resuspended in 180 μL M9 worm buffer (final volume 200 μL/worm) before transfer to 96-well plates for serial dilution in 1X phospho-buffered saline (PBS).

For 96-well plate disruption, bleached worms were rinsed 1X in M9 worm buffer + 0.1% Triton X-100 to remove bleach. Worms were then treated for 20 minutes with 100 μL of a SDS/DTT solution in M9 worm buffer (0.25% sodium dodecyl sulfate + 3% freshly mixed dithiothrietol, chemicals from Sigma Aldrich), rinsed with 150 μL M9 worm buffer, then rinsed again in 1 mL M9 worm buffer + 0.1% Triton X-100 before transfer to 3 mL M9 worm buffer + 1% Triton X-100 in a small (40 cm) petri dish (Fisher Scientific). A deep-well plate (2 mL square well plate, Axygen) was prepared by addition of a small quantity of sterile autoclaved 36-mesh silicon carbide grit (Kramer Industries) to each well, followed by addition of 180 μL M9 worm buffer + 1% Triton X-100. Individual worms were then pipetted out in 20 μL aliquots of buffer and transferred to individual wells of the plate. The plate was covered with parafilm and allowed to chill for at least one hour prior to disruption to reduce heat damage to bacteria. Parafilmed plates were capped with 96-well square silicon sealing mats (Axygen Impermamat AxyMat Sealing Mats) and disrupted by shaking in a 96-well plate shaker (MO-BIO Laboratories, shaking at 30 hz for 1.5 minutes before flipping plate and shaking an additional 1.5 minutes to ensure even disruption). Plates were then centrifuged at 2500 x g for 2 minutes to collect all material in the bottom of the wells, then all plate contents were resuspended by pipetting and transferred to 96-well plates for serial dilution in PBS.

### Bimodality

Bimodality was assessed using the bimodality coefficient [54]:

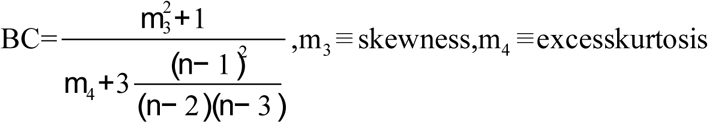

with the canonical value of BC_crit_ = 0.55 used to mark the transition from a unimodal to a bimodal distribution. Calculations were implemented in Python using scipy.stats.

### Parameter estimation

*Growth rate and carrying capacity*. Birth rate *b* and carrying capacity *K* were estimated simultaneously by fitting a logistic growth model to data for growth of bacteria inside the worm (**Fig. 2**). Briefly, *C. elegans* AU37 was fed on live *E. coli* MC4100 at ~10^9^ CFU/mL for two days, then washed 1X with M9 worm buffer + 0.1% Triton X-100 and 2X with M9 worm buffer to remove most external bacteria, then resuspended in 1 mL S medium + 2X heat-killed OP50 in a 24-well plate, covered with an adhesive gas-permeable microplate sealing film, and incubated at 25°C for 10 days. To minimize new colonization, worms were washed and re-fed in fresh S medium + 2X heat-killed *E. coli* OP50 at 24-hour intervals, and 12 worms were surface bleached and individually digested for CFU plating.

Maximum growth rate was estimated by allowing *E. coli* MC4100 bacteria to grow outside the worm in S medium + 2X heat-killed OP50 at 25°C - essentially identical to culture conditions used in worm assays, but without the worm environment. Due to the high turbidity of the medium with heat-killed OP50, CFU counts over time were determined via dilution plating.

*Net growth rate*. From the data in **Figure 2**, we observed that the net growth rate (births-deaths) of *E. coli* MC4100 inside the worm intestine was very low, ~0.06 hr^-1^; bootstrap resampling and refitting of the logistic equation (10,000 runs) produced a 95% CI of 0.21 ≥ b-d ≥ 0.029 hr^-1^. This low net growth rate implies that birth and death rates inside the worm are very close to one another. The existence of zero counts (empty worms) in the first days of outgrowth implies that stochastic extinction in lightly-colonized worms occurs at an observable frequency, consistent with expectations for small populations where birth is only slightly more likely than death. (We must also note that the lack of uncolonized worms at later time points implies that some colonization is occurring during the outgrowth period, due to growth of excreted bacteria in the liquid environment in the 24 hours between washes.)

As it is very difficult to estimate birth rate inside the worm in isolation, we are forced to extrapolate. The net growth rate (*b-d*) should be small, and as stated in the main text, there is a large amount of variation in bacterial population size between worms; in the context of a stochastic simulation, we therefore choose values of *b* and *d* which are as large as possible given the physiological constraints of the system in order to maximize stochastic noise. For simulations, we use the estimated growth rate of *E. coli* MC4100 outside the worms in these growth conditions (~0.6 hr^-1^) as a maximum birth rate and choose a death rate of 0.54 hr^-1^ to produce a net growth rate of 0.06 hr^-1^.

*Excretion and death rates*. Total death and excretion were estimated using a chloramphenicol exposure assay, where chloramphenicol at concentrations above the MIC for this bacterial strain are used to prevent bacterial division without causing cell death. Here, change in CFU/worm over time is used to estimate total death, and change in CFU/mL outside the worms is used to estimate excretion (which in our model is incorporated into the cell death term *d*).

Briefly, worms were colonized with *E. coli* MC4100 E3350 or E3322 at ~10^9^ CFU/mL for 2 days, then rinsed and purged of non-attached bacteria as previously described. Worms were then washed 1X in M9 worm buffer + 0.1% Triton X-100, surface bleached as previously described, and moved to 1 mL S medium + 0.2X heat-killed OP50 + chloramphenicol at a range of concentrations (0, 10, 20, 50, 100, and 200μg/mL) in a 24 well plate covered with an adhesive gas-permeable microplate sealing film. The plate was incubated with shaking at 300 RPM at 25°C for 48 hours; samples of supernatant and batch worm digests (160-230 worms/digest) were taken for all conditions at 0, 24, and 48 hours.

Excretion was estimated from CFU/mL counts outside the worms at 24 hours over this range of chloramphenicol concentrations (**Fig. S1A**). From these data, excretion rates are approximately 1 CFU/10^3^-10^4^ intestinal CFU/hr. Excretion therefore appears to account for only a small fraction of total cell death in intestinal bacteria in the worm.

Death rates were estimated from CFU/worm measurements over time (**Fig. S1B**). At high chloramphenicol concentrations (50-200 μg/mL), change in CFU/worm between 24 and 48 hours after exposure was negative and effectively constant across drug concentrations (**Fig. S1B**), indicating that the antibiotic was effectively suppressing growth of bacteria inside the worms. These data were used to estimate total average death rate of bacteria inside the worms as 0.05-0.1 hr^-1^. If we consider the experimentally derived estimate of death rate (0.05-0.1 hr^-1^) to be accurate, this would imply a very low birth rate of 0.1-0.15 hr^-1^.

*Colonization rate*. Colonization was estimated using a short-time feeding assay. Briefly, *C. elegans* AU37 was fed on live *E. coli* MC4100 at a range of concentrations from ~10^9^ to 10^6^ CFU/mL (corresponding to colonization rates of 0.1 - 100 CFU/worm/hr). Bacterial density was quantified by dilution plating. At 1 and 4 hours after feeding, 30-60 worms were removed from culture, washed and surface bleached as previously described, and disrupted in a batch; average CFU/worm was determined by dilution plating. The experiment was performed twice, and data were pooled for parameter estimation (**Fig. 2**).

*Feeding rate*. Feeding rate was estimated from the same short-time feeding assay by comparing CFU/mL outside the worms at time 0 and after 10 hours of feeding. Depending on initial concentration, 5-60% of total bacteria in each well were consumed over this time. The resulting ΔCFU/mL was divided by the total number of worms in each well to get an estimated average CFU eaten/worm/hr (**Fig. 2**).

*Robustness to parameter choice*. The transition from bimodal to unimodal population distribution with increasing colonization rate is robust to the parameter values used in simulation (**Fig. S8**). Holding (*b-d*) constant, we can observe that this transition occurs around *c* / b=1 across a range of parameter values (compare [*d*=2.44, *b*=2.5] with the other parameter sets). The observed transition is therefore robust to the uncertainty in the parameter values extracted from experimental data (compare [*d*=0.1, *b*=0.16] and [*d*=0.54, *b*=0.6]). However, the variation in total number of bacteria across individual simulated worms increases dramatically as the values of (*b,d*) increase, as does the frequency of extinctions; this is expected behavior for a model where death is stochastic.

### Microscopy

After surface bleaching and rinsing as previously described, worms were paralyzed in M9 worm buffer with 10% sodium azide for 10 minutes before transferring to slides with 2% agarose pads for microscopy. Microscopy was performed on a Nikon Eclipse T*i* inverted light microscope system with Chroma filter sets ET-dsRed (49005), ET-YFP (49003), and ET-CFP (49001) and a Pixis 1024 CCD camera; MetaMorph microscopy automation and image analysis software (Molecular Devices) was used for machine control and image capture and integration.

### Stochastic modeling

Gillespie stochastic simulation of the logistic equation-based stochastic model was performed using the PySCeS environment (Python Simulator for Cellular Systems, http://pysces.sourceforge.net) and StochPy (Stochastic modeling in Python, http://stochpy.sourceforge.net) in iPython Notebook. The tau-leaping method was used in these simulations to reduce computation time. The model used in these simulations (neutralLV.psc) is presented:

NeutralLV.psc

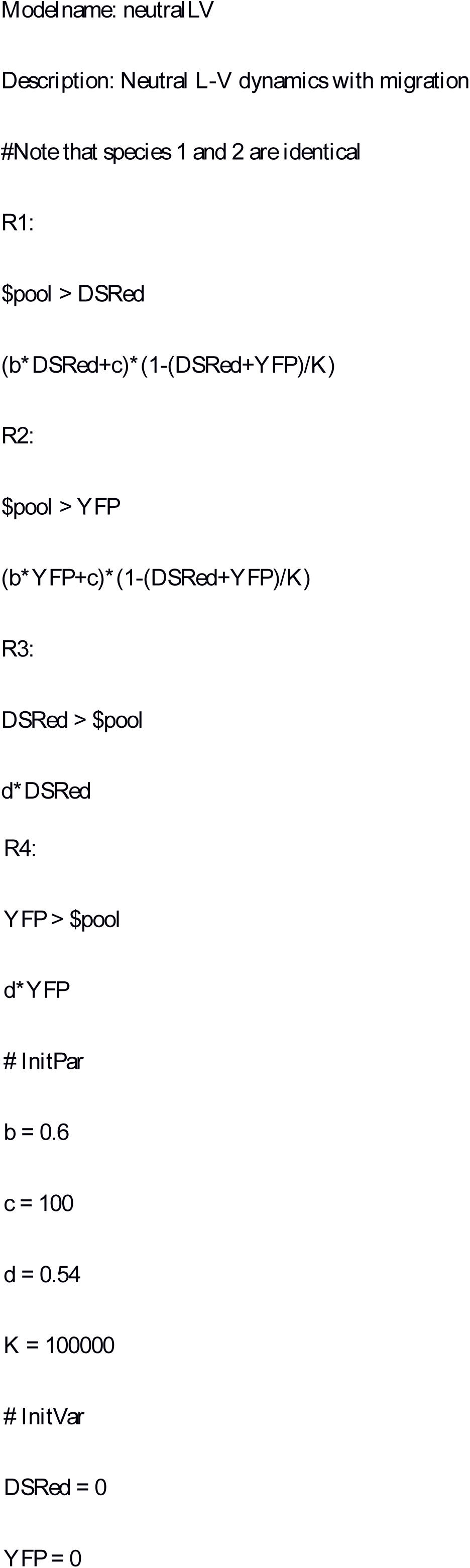

## Acknowledgements

We thank the current and former graduate students and postdocs of the Gore lab, especially J. Friedman, T. Biancalani, and K. Korolev, for discussion that improved this manuscript. Special thanks to K. Axelrod and S. Gandhi for fueling our efforts.

